# The blind choreographer: evolution of social norms and correlated equilibria

**DOI:** 10.1101/435586

**Authors:** Bryce Morsky, Erol Akçay

## Abstract

Social norms regulate and coordinate most aspects of human social life, yet they emerge and change as a result of individual behaviours, beliefs, and expectations. A satisfactory account for the evolutionary dynamics of social norms therefore has to link individual beliefs and expectations to population-level dynamics, where individual norms change according to their consequences for individuals. Here we present a new model of evolutionary dynamics of social norms that encompasses this objective and addresses the emergence of social norms. In this model, a norm is a set of behavioural prescriptions and a set of environmental descriptions that describe the expected behaviours of those with whom the norm holder will interact. These prescriptions and descriptions are functions of exogenous environmental events. These events have no intrinsic meaning or effect on the payoffs to individuals, yet beliefs/- superstitions regarding them can effectuate coordination. Though a norm's prescriptions and descriptions are dependent upon one another, we show how they emerge from random accumulations of beliefs. We categorize the space of social norms into several natural classes and study the evolutionary competition between these classes of norms. We apply our model to the Game of Chicken and the Nash Bargaining Game. Further, we show how the space of norms and evolutionary stability is dependent upon the correlation structure of the environment, and under which such correlation structures social dilemmas can be ameliorated or exacerbated.

---

Cooperation and coordination in social settings are fundamental to the functioning of human societies. A simplistic view of human behavior as purely self-interested actors would suggest that cooperation and coordinated action should be rare if there are conflicts of interests between players and no external incentives (or coercion) is at hand to resolve these conflicts. And yet, we observe many examples of cooperation without external incentives, mediated by informal mechanisms such as social norms [29, 14]. More generally, it is apparent that while humans do tend to make decisions to improve their payoffs, individual decision making is also imbued with both prescriptive beliefs on their own behavior and expectations on others’ [16]. These prescriptions and expectations may or may not be rooted in reality or accurately describe others’ behaviors, but regardless have a real effect on human behavior [24, 10, 13]. Importantly, they can induce regularities in a population that allow self-interested agents to cooperate and coordinate with each other efficiently.

Here, we focus on social norms, defined as a set of informal behavioural rules that help govern human interactions on interpersonal to societal levels [5]. Social norms can be both descriptive and prescriptive. Descriptive norms tell us what behaviours to expect from others, whereas prescriptive norms tell us how to behave and can be predicated on descriptive norms [9]. Although norms coordinate behaviour, they generally are not explicitly designed by individuals or society.

Social norms that facilitate cooperation are commonly modeled through the perspective of game theory [40, 18, 30]. One can model norms as mechanisms selecting amongst (the typically numerous) Nash equilibria in repeated games [7]. Alternatively, social norms can consist of conditional behaviour rules [5] that convert social dilemmas into coordination games, for example by external enforcement (either through institutions or by others in a group) [8, 37]. Our focus is on social norms that lack external enforcement, but still are followed because agents, given their beliefs about the world (also induced by social norms), find it in their interest to do so. More specifically, we model social norms that act as “choreographers” [17] that induce correlated beliefs in agents allowing them to coordinate on a *correlated equilibrium* [2, 3] of the game.

A correlated equilibrium is generated by a “correlating device” (the “choreographer”) that suggests pure strategies to the players. If the suggestions to each player are correlated in a certain way, and the agents know these correlations, it can be optimal for the players to follow the suggestions of the choreographer. Such behavior is called a correlated equilibrium. The Game of Chicken is an excellent exemplar for the coordinating effects of correlated equilibria as it can feature a scenario in which coordination would lead to a socially optimal yet game-theoretically unstable state. Consider an intersection that two cars approach from perpendicular directions. The drivers may choose to drive through the intersection or stop. The payoff matrix for this game is

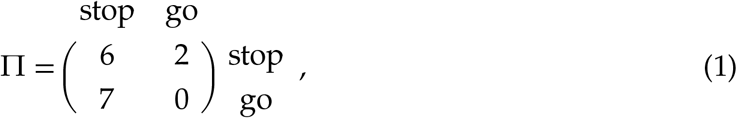

where *π*_*ij*_ ∈ Π is the payoff to a player playing the row strategy vs an opponent playing the column strategy. The cooperative strategy is to stop, which is to “chicken out,” and both players stopping is the socially optimal state. However, at the *Nash equilibrium,* the frequency of cheating (i.e. going) is ⅓. Now imagine a set of traffic lights at the intersection. If these traffic lights only ever both signal stop, or one signals go and the other stop, then a crash can be prevented. In fact, if the probabilities of these signal pairings are chosen correctly (such as probability ⅓ for each of the pairings), it is rational for the drivers to obey them given knowledge of how the lights operate, and we thus have a correlated equilibrium.

Correlated equilibria may emerge from learning and other adaptive procedures [15, 21, 20, 23, 1], and can be viewed as an outcome of evolution [11]. In the classical conception of correlated equilibrium, there is a third party that decides on the correlations between the suggestions. Here, we take out this third party and ask how correlated strategies arise and compete against each other decentrally. In other words, we ask whether correlated equilibria can evolve in a world where people do not have to agree on the correlating device and there is no external enforcement.

To answer this question, we model a world where nature provides a collection of events with no inherent impact on individuals, but possessing correlations amongst themselves. We propose that individuals might choose to interpret these events normatively as prescriptions on their own behaviors and descriptions of the expected behavior of others. Such “superstitious” beliefs on the meaning of events in nature can help individuals exploit the pre-existing correlations as coordinating devices. Returning to our traffic light analogy, we consider events that are potential signals at the intersection. We show how belief structures about these events can emerge, and be both individually rational and evolutionarily stable even though it is irrational to observe any meaning behind the events. We are particularly interested in this phenomenon with respect to non-cooperative games that feature a social dilemma where the socially optimal solution is evolutionarily unstable. As illustrative examples, we apply our general framework to the Game of Chicken and the Nash Bargaining Game.

## 1 The model

Consider two players meeting to play a game. Each player privately observes an event, which is correlated with their opponent’s event and other events in ways unknown to the players. Each of these events can be assigned a label by which players can differentiate them. However, the events and labels inherently have neither meaning nor an association with one another [12]. As a function of the observed label, players’ social norms prescribe a pure strategy recommendation and also describe (to the focal player) the expected behaviour of the opponent. The players then may obey the norm’s prescription or play a default strategy from which they will earn a payoff. Though players cannot observe the prescriptions their opponent’s observe, they know the conditional probability distribution of prescriptions their opponent may be receiving given the recommendation they themselves have been given. A social norm then is the set of these labels, prescriptions, and descriptions; which are obeyed if they form a correlated equilibrium. The key to obeyance of the norm is that ignorance of their opponent’s exact behaviour permits a different fitness than the Nash equilibrium.

Mathematically, we model the world of events as an undirected graph, *G* = (*V,E*), where the edges, {υ_*i*_,υ_*j*_} ∈ *E*, represent an interaction between two players via a game and the vertices privately observed events. We assume all the games players play are the same and the private events have no direct effect on the payoffs. Players then play each other a large number of times, such that they observe all events. When a game is played, each player is equally likely to be privately assigned either vertex of the edge. In the event that the edge is a self-loop, both players are assigned the same vertex (though this is still privately observed). The probability of selecting edge {υ_*i*_,υ_*j*_} is *a*_*ij*_ ∈ *A*, a weighted symmetric adjacency matrix. We limit the number of ways by which a player can distinguish vertices by defining a set of labels, *L*, to apply to each vertex. If two vertices have the same label, then the player cannot differentiate between the two. In matrix notation, we define this as follows. Given a set of labels *L,* let *L* be the |*L*| × |*V*| *label matrix*, which partitions the vertices into |*L*| labels (i.e. *l*_*ij*_ = 1 if *υ*_*j*_ has label *l*_*i*_, otherwise *l*_*ij*_ = 0). Norms prescribe pure strategy recommendations to labels. We can represent the behaviours players are prescribed by the |*S*| × |*L*| *prescriptive behaviour matrix, P*, where *p*_*ij*_ = 1 if the norm prescribes the player to play *s*_*i*_ given *l*_*j*_. If |*L*| = |*V*|, players can be prescribed a particular strategy for each vertex individually. In the case where the norm has no prescription for a label, the players play a default strategy, a symmetric (mixed) Nash equilibrium for the game. The expected behaviour of an opponent is determined by the column stochastic |*S*| × |*L*| *descriptive behaviour matrix, D*, which is the expected behaviour of an opponent given a label observed by the focal player. Given these constructions, a norm is defined by the matrix triplet (*L, P, D*).

Figure 1 details our model pictorially for a four event graph with two players having differing norms. An event is chosen where each player observes a cat. In this case, player 1 observes the white cat and player 2 the black cat. Player 1’s norm prescribes cooperation when it observes black or white cats, and defection for grey and tabby ones. On the descriptive side, Player 1’s norm specifies a conditional distribution of behaviours for Player 2. We assume that this distribution is conditional on the labels of the focal player. In other words, Player 1’s expectation of Player 2’s behaviour is the same when observing black and white cats. In the example in Figure 1, Player 1 believes that Player 2 will cooperate with probability ¼ and defect otherwise. Notice that the expectations created by a player’s norm can be different than the actual distribution of prescriptions received by the opponent. This discord is illustrated by player 2’s norm, which could well be described as impudent; they believe they should defect regardless of which cat they see and expect their opponent to always cooperate.

**Figure 1:**
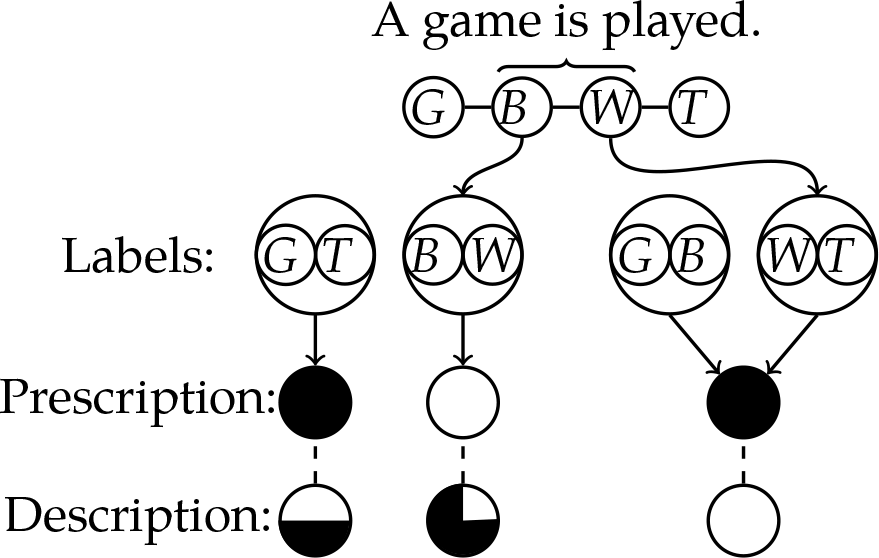
A pictorial representation of the model. Here, each vertex on the graph corresponds to the observation of a different colored cat (G: gray, B: black, W: white, T: tabby). As an example, Player 1 and 2 meet and they observe a black (B) and white (W) cat, respectively. Player 1’s norm recommends that they cooperate (represented by the white), and — given that player 1 has been told to cooperate — claims that player 2 will cooperate with probability ¼. Player 2’s norm recommends they defect (represented by black), and claims that player 1 will always cooperate.

We examine the effects of a variety of competing norms on a given event graph. In playing an opponent with a differing norm, a player will still follow its own norm and thus not necessarily receive the expected payoff their norm would dictate. Norm change can be facilitated by such disparity between expectations generated by descriptive norms and reality [6]. However, we assume that players do not reflect upon — or at least do not do so as to modify — their norms directly, but can observe the fitness of competing norms. Thus, norm change is generated by imitation dynamics, whereby players imitate the norms of those with higher fitness [22]. The expected payoff of following norm *n* employing *P* playing norm *n*’ with *P*’ is a function of the prescriptions for each player at each interaction multiplied by the payoff from that strategy pairing. Mathematically, this is

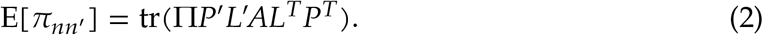

Fitness is thus the sum of this expected payoff over all norms weighted by their frequency.

## 2 Results

### 2.1 A classification of norms

Our definition of social norms captures the idea that norms imbue an inherently meaningless world with prescriptions and expectations about self and others’ behaviors. However, we do not assume the prescriptions of a social norm are to be blindly obeyed. In particular, we assume that a norm will only be obeyed when — given the expectations created by the descriptive aspect of the norm — individuals cannot improve their payoffs by deviating from the social norm’s prescriptions. In this section, we will first define the rationality condition and classify important norms. Figure 2 depicts a Venn diagram of our classification of norms (a summary of this classification can also be found in the appendix, Table 1). In section 2.2, we will address evolutionary concerns.

**Table 1:** Summary definitions of norms and concepts.

| Social norm /<br>concept | Definition | Mathematical representation |
| --- | --- | --- |
| Null norm | A default strategy (mixed Nash) is played, since it is irrational for the norm holder to obey the norm. | $\text{tr}(\Pi D(P - \tilde{P})^T) < 0$. |
| Rational norm | The norm's prescriptions are played, since it is rational to obey the norm. | $\text{tr}(\Pi D(P - \tilde{P})^T) \geq 0$. |
| Empirically<br>validatable norm | There exists a rational norm whose prescriptions are such that they are consistent with the focal norm's descriptions. | $\exists n'$ such that $n'$ is rational and $D = P' L' A L^T$. |
| Consistent norm | A norm that is empirically validatable with itself. | $n$ is empirically validatable and $D = P L A L^T$. |
| Inconsistent norm | A norm that is empirically validatable with a different norm and not itself. | $n$ is empirically validatable and $D \neq P L A L^T$. |
| Correlated<br>equilibrium | Two norms form a CE if they are empirically validated by one another. | See Appendix A. |
| Evolutionary stable<br>norm | A norm that is evolutionarily stable. | $\forall n' \neq n, \text{tr}(\Pi P L A(P L - P' L')^T) > 0$, or<br>$\text{tr}(\Pi P L A(P L - P' L')^T) = 0$ and<br>$\text{tr}(\Pi P' L' A(P L - P' L')^T) > 0$. |
| Best response norm | The prescriptions at each edge of the graph are best responses to one another. | $\forall \{v_i, v_j\}$, the prescription at $v_i$ is the best response to the prescription at $v_j$. |

**Figure 2:**
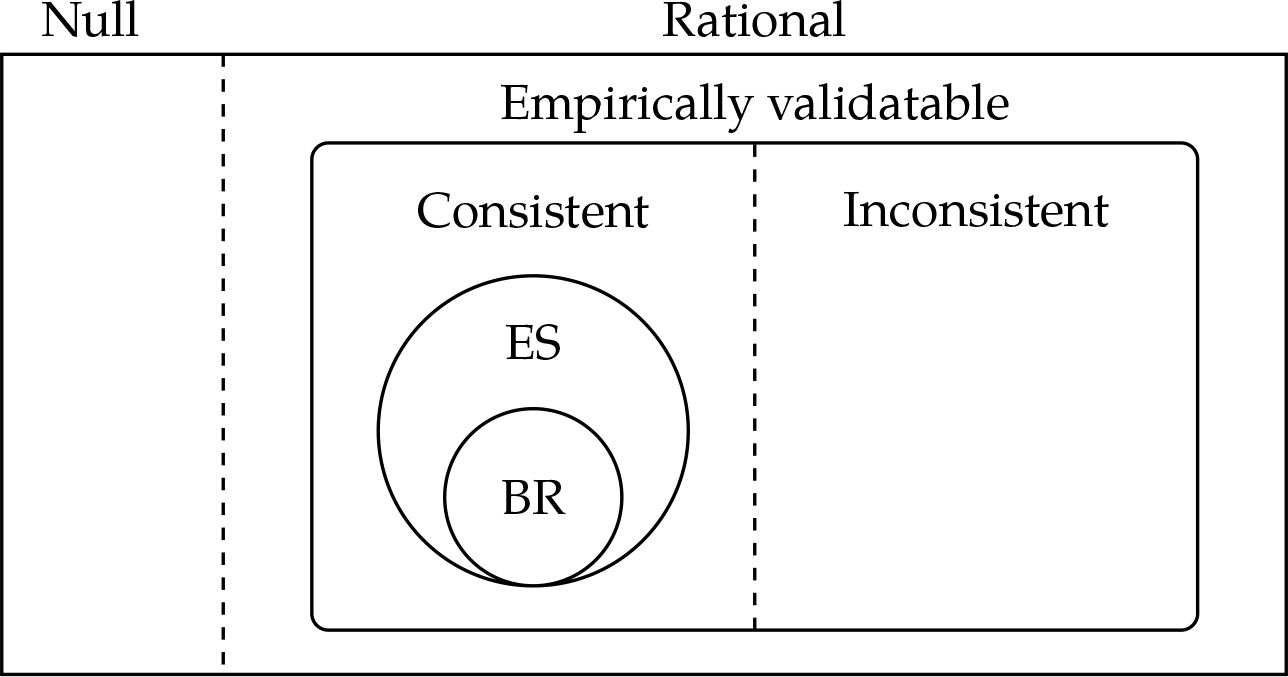
A Venn diagram of the set of norms. All norms can be partitioned into null and rational norms. Empirical norms can be partitioned into consistent (which implement a correlated equilibrium when playing itself) and inconsistent norms. Inconsistent norms may implement a correlated equilibrium with complementary inconsistent norms, but not with themselves. Best response norms (BR) are a subset of evolutionary stable norms (ES). A summary of these norms and concepts are found in Table 1.

Let (*L, P, D*) be a social norm. The expected payoff for obeying the prescription at label, *l*_*i*_ is (Π*D*)_\**i*_ · *p*_\**i*_. It is rational for a player to obey their norm’s prescriptions if each prescription at a label is a best response to the expectation of an opponent’s behaviour (descriptive norm) at that label, i.e.

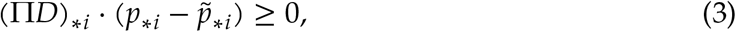

for every label *l*_*i*_ and prescription p̃_\**i*_ ≠ *p*_\**i*_. If a norm does not obey condition 3, then players who have it will disregard the recommended prescriptions and play a default strategy at all events. We will call such norms *null norms*. If the game features a dominant strategy, we assume that this is the default. However, if the game features a symmetric mixed Nash equilibrium, we assume that that is the default. In the case of multiple such strategies, we have a family of null norms.

Although the descriptive aspect of a norm does not have to accurately describe opponent behavior, there exists a class of norms that does fit the behavior of (possibly hypothetical) opponents. In particular, a norm is *empirically validatable* if its descriptions match the behaviour of a rational norm. This class is of interest, since two norms that are empirically validatable with respect to one another implement a correlated equilibrium when they play one another (see appendix A for a proof).

We are also interested in a particular class of norms where everyone would obey their norm given that everyone else also obeys the same norm [4]. We call these norms *consistent norms*. In other words, the descriptive part of a consistent norm accurately describes the behavior of opponents that follow the prescriptions of the same norm, given the underlying event graph. In contrast, the descriptive part of an *inconsistent norm* does not follow from its own prescriptive part, but is rather conditioned on a player playing a different norm. For example, consider the event pair {black cat, white cat} in which the two cats are completely correlated with one another. Suppose, when a black cat is observed, a norm prescribes defection and expects cooperation from the opponent. If this norm is consistent, it will prescribe cooperation and expect defection when a white cat is observed. On the other hand, an inconsistent norm can hypocritically prescribe its carrier to also defect and expect cooperation in the event the carrier sees a white cat. Mathematically, if a norm is empirically validatable and *D* = *PLAL*^*T*^, then the social norm is consistent. Note that if a norm is consistent, it necessarily implements a correlated equilibrium playing against itself. However, if a norm is empirically validatable but *D* ≠ *PLAL*^*T*^, then the norm is inconsistent. The left hand norm in Figure 1 is consistent and the right inconsistent. It is worth reminding that players do not explicitly know the event graph, and therefore cannot directly evaluate whether the norm is consistent or inconsistent. Nonetheless, as we will see below, consistent and inconsistent norms have fundamentally different evolutionary properties.

We label a subset of consistent norms where the prescriptions for each vertex of an edge are best responses to one another as *best response norms*. Best response norms always exists if the game admits a single strategy Nash equilibrium. However, for a 2 strategy pure Nash equilibrium, a best response norm exists if and only if the graph is bipartite. Since, we may assign to each partition one of the strategies and thus each vertex along an edge is a best response to each other. The fitness of a best response norm playing itself is the mean of the payoffs of the pure Nash equilibria.

### 2.2 Evolutionary stability of norms

Similar to the concept of an evolutionary stable strategy [35], an *evolutionary stable norm* is a norm that cannot be invaded by an initially rare norm (see appendix B). We are concerned with the evolutionary history initiated at a state without prescriptions where everyone plays a null norm. As long as the prescriptions are not obeyed (3 is not satisfied), they may increase in the population by drift. However, upon being rational, they can be obeyed. If such a norm is a consistent norm, it may then invade if the null norm plays a mixed Nash strategy. Since, any strategy within the support of a mixed Nash — which all consistent norms are, as they implement a correlated equilibrium — playing the mixed Nash strategy receives the mixed Nash payoff. Further, as a consistent norm implements a correlated equilibrium, its payoff playing itself is greater than any alternative norm under the same event labeling. Since the null norm’s behaviour is identical across all events, the labeling is inconsequential, and thus receives a lower fitness playing the consistent norm. Therefore, the consistent norm can invade. More generally, the same arguments applied to a subset of events show the emergence of obeyed prescriptions from null prescriptions. In other words, the evolutionary dynamics of norms tend to manufacture beliefs that coordinate interactions out of inherently meaningless events.

Continuing with the assumption that the null norm plays a mixed Nash, evolutionarily stable norms must be consistent norms. Since consistent norms induce a correlated equilibrium, they can be evolutionarily stable against norms that share the same relative labeling of events (i.e. so long as events are labeled such that they are differentiated in the same ways) [26]. However, norms with different labels may invade, unless the resident norm is a best response norm. Unlike consistent norms, inconsistent norms cannot be evolutionary stable, because they can be invaded by a norm that validates their descriptions, which by definition is different from themselves. Therefore, we may have a polymorphism of two inconsistent norms if they empirically validate one another. If the successful invader is inconsistent but the norms do not empirically validate one another, the situation is more complicated and we may have a polymorphism or dominance by the invader. However, if the successful invader of an inconsistent norm is a consistent norm, then the consistent norm is evolutionarily dominant with respect to the inconsistent norm.

Polymorphisms of weakly evolutionarily stable consistent norms with differing prescriptions can exist so long as on average the pairings of prescriptions are the same, or, trivially, if the null norm and consistent norms’ behaviours are identical. This latter condition occurs if the game has only a single strictly dominant pure Nash equilibrium. Since, the only correlated equilibrium is then to always play the pure Nash strategy and thus all consistent norms must prescribe only it. If there is a single non-dominant Nash equilibrium, this is not true in general (see appendix C for an example). Regardless, we are primarily interested in games where there are several pure Nash equilibria and a mixed Nash equilibrium, since it is these cases in which a social norm can be used to coordinate behaviour.

### 2.3 Examples

#### 2.3.1 The Game of Chicken

The existence and stability of norms for a game is dependent upon the underlying event space. Consider the bipartite graph 3A for the Game of Chicken. There is one behaviour that a null norm can play, namely the mixed Nash equilibrium where they cooperate at each vertex with probability ⅔. The set of consistent norms is composed of five norms (*n*_1_, *n*_2_, *n*_3_, and the inverses of *n*_2_ and *n*_3_), three of which are stable (norms *n*_1_, *n*_2_, and the inverse of *n*_2_).

Continuing with the example above, we may consider all prescription as a strategy from which we have a 12 dimensional replicator equation, the symmetric Nash equilibia of which are the fixed points [22]. There are 25 fixed points of this model, 3 of which are stable (the monomorphic states of the three stable consistent norms). For monomorphic populations, *n_1_* has a higher fitness than the null norm and the two best response norms, *n*_2_ and its inverse. Contrast this to the graph 3B also playing the Game of Chicken, which is also a bipartite graph. The evolutionary stable norms of graph 3B are all best response norms, which have a lower monomorphic fitness than *n*_1_.

#### 2.3.2 The evolution of fairness in the Nash Bargaining Game

The Nash Bargaining Game provides us with another interesting example [28]. Each player’s strategy is a demand of a portion of a dollar. If the sum of the demands is less than or equal to a dollar, then each player receives their demand. If, however, the sum exceeds a dollar, then they both receive nothing. A variety of solutions (corresponding to varying definitions of fairness) have been found [28, 25, 19]. We are interested in how a 50-50 division of the dollar can emerge. It has been shown that a single population replicator equation model can converge to such a fair distribution for a symmetric linear Nash Bargaining Game [34]. However, the history of play matters, and the basin of attraction to a suboptimal polymorphic stable equilibrium is not negligibly small. In a model of tenants and landlords playing the Nash bargaining game, a fair division of the surplus between tenants and landlords is stochastically stable if memory is large and noise small [42]. Here, we will have our players play the linear symmetric Nash Bargaining Game at each edge. Further, we admit only a finite number of strategies that divide the dollar (see Appendix D for details).

Let us begin with a discussion of the complete graph with more than one vertex such as Figure 3C. We are considering monotonically competitive process, whereby a mutant may replace the resident in an evolutionary chain. Consider consistent norms that assign *δ* ≤ ½ to themselves at every vertex but for one at which they assign 1 − *δ* to themselves.

**Figure 3:**
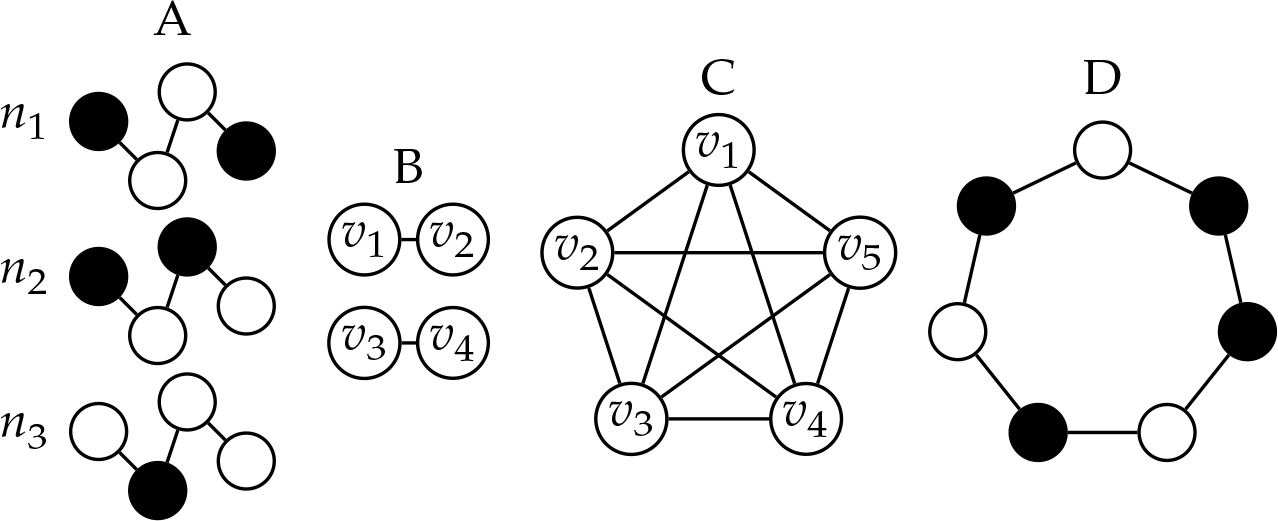
Several interesting event space graphs. White corresponds to cooperate and black to defect in graph *A*. *n*_1_ and *n*_2_ are evolutionary stable norms. *n*_1_ provides a higher fitness than *n*_2_, which is a best response norm. Black corresponds to demanding *5* and white to demanding 1 − *δ* in graph *D* for the Nash Bargaining Game.

Such a norm can be vulnerable to a mutant assigning ½ ≥ *δ*′ > *δ*. Therefore, we find that a norm employing

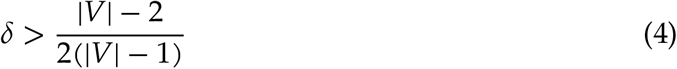

is a monomorphic evolutionary stable norm. The space of monomorphic evolutionary stable norms decreases as we increase the number of vertices, and thus we approach the fair distribution being the only monomorphic evolutionary stable norm. Since each event is equally correlated with every other event, there is a *Veil of Ignorance*, with respect to every vertex, leading to fairness [32]. In contrast to the complete graph, the cycle graph with degree 2 can display suboptimal solutions and varying degrees of fairness across the graph. If the graph is odd, the only best response norm is *δ* = ½, yet any complete division of the dollar is a best response norm for an even graph. Additionally, we may have suboptimal evolutionary stable norms as depicted in Figure 3D.

We explored evolution of norms for the complete graph, ring graph with degree 2*K*, and small world graphs [41] via trait substitution simulations (details in Appendix D). These simulations permit polymorphisms of inconsistent norms (depicted in black). Note that the monomorphic states are monomorphic with respect to behaviour and not labels; regardless of the label one attaches to a vertex, we are interested in the behaviours and outcomes. Figure 4 depicts the results for the complete and *K* = 1 ring graph (remaining results are in the appendix). Although the fair distribution is frequently reached, polymorphisms of inconsistent norms can exist and lower the mean fitness. For example, at *α* = 4 on the complete graph, we have a third of the population demanding ¼ of the dollar and the remaining demanding ¾, from which we have a mean fitness of ¼ (panel **e**). The ring graph (**c**) has much more diversity than the complete graph (**a**). Increasing *α* decreases the frequency of the fair outcome. For the complete graph (**b**), this generates a greater diversity of polymorphisms (most having higher fitness, though some lower). Increasing *α* on the ring graph decreases the diversity of polymorphisms as can be seen in the defined bands of panel **d**.

**Figure 4:**
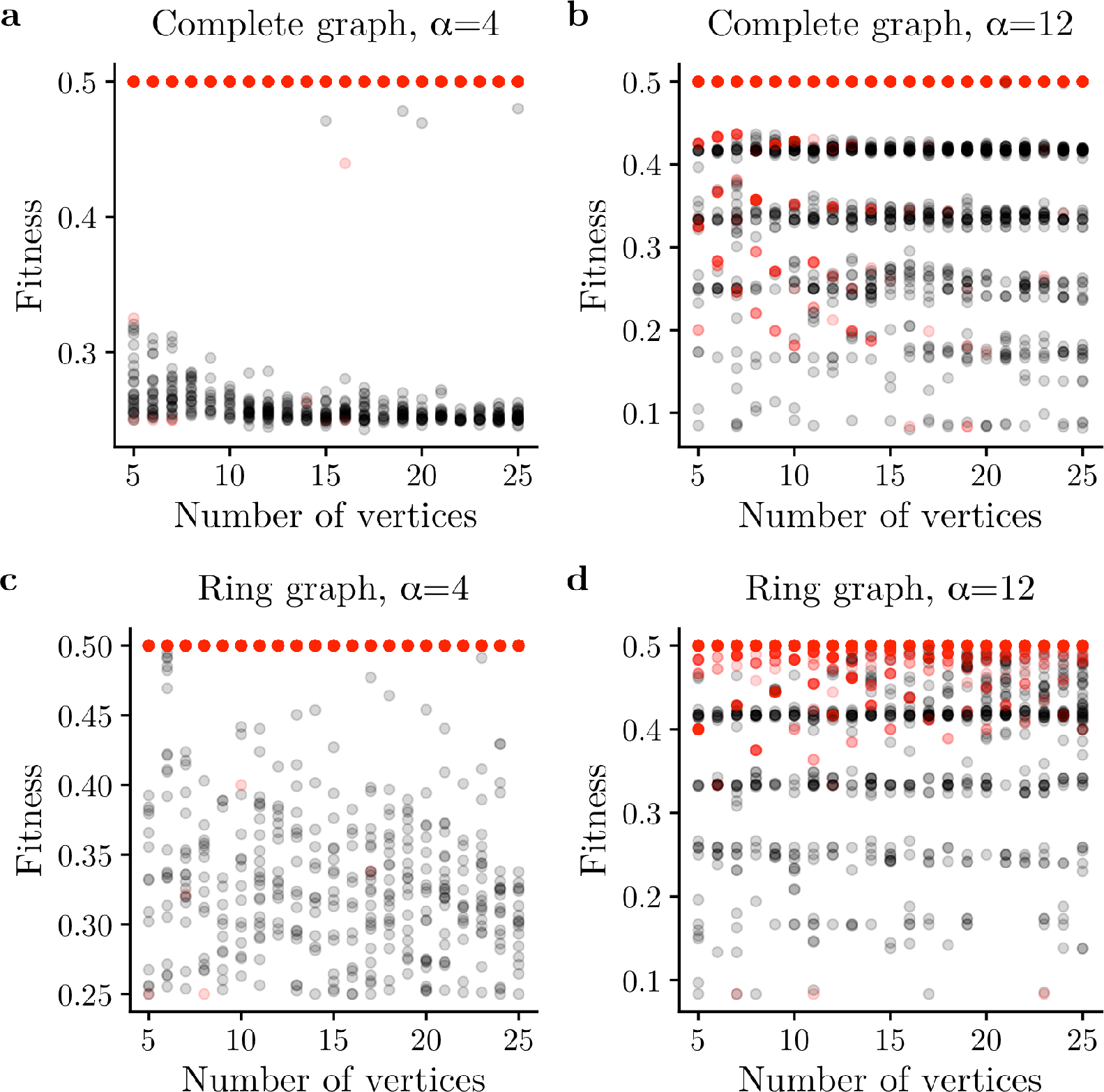
Mean fitness frequency in the Nash Bargaining Game for the complete and ring graphs, and for dollar refinements *α* = 4,12. Red corresponds to monomorphic states, and black polymorphic.

## 3 Discussion

In this paper, we show how social norms that coordinate behaviours can be bootstrapped from random superstitions about irrelevant events in the world. Our model of the evolution of correlated equilibria differs from previous work in that we model a completely decentralized scenario with no commonly agreed correlating device. For instance, in [27], new signals arise through proposals that all agents have to agree to (i.e., new signals must increase all agents’ payoffs). [26] consider a case that is closer to our setup, where a given correlating device recommends behaviours, but individuals may choose to not follow recommendations. They define *evolutionarily stable correlation* as a correlated equilibrium which is evolutionarily stable with respect to mutants that disobey the signals given. However, unlike our model, [26] do not consider the evolution of different correlating devices through selection on individuals, and do not require responses to recommendations to be individually rational given expectations.

Another key to our model is the synthesis of prescriptive and descriptive norms. Inconsistencies between between prescriptive and descriptive norms have been experimentally shown to weaken norm adherence [31,36]. As such, our rationality and empirical validation conditions are key to aligning prescriptive and descriptive norms and inducing their obeyance. We show that empirically validatable norms not only align the internal norms of players, but also may align their behaviours by inducing a correlated equilibrium. Such social norms act as choreographers of the players’ behaviours potentially ameliorating social dilemmas, a well studied product of social norms [39]. However, our players and the choreographer are blind to the true correlations between events in the world, and as such this coordination is unintentional. Rather than being designed, choreography can emerge from the accumulation of random superstitions of the environment. The natural cues in the environment upon which the norms’ prescriptions are based, whether they be sight of a coloured cat or something equally superstitious, have no inherent meaning. Rather, the meaning of events in terms of actions to be taken are invented by individuals themselves, and together with the beliefs about others’ behaviors, can spread through the population through imitation.

In sum, we show that normative meaning of random events in nature can emerge in a population. In alignment with empirical work [33, 38], our norms begin individually, but through internal conditions, competition, and an imitation process become standardized and stabilize. The null norm is weakly unstable when the default behaviour is a mixed Nash, and thus forms a basis from which mutations in the norm can induce them to be followed. Within the set of these rational norms are those that are empirically validatable, and within that set are norms that induce a correlated equilbria. In this way, players can coordinate their play through decentralized evolution of arbitrary meaning.

## Appendix A Correlated equilibria

Here we show that two norms that empirically validate one another form a correlated equilibrium. To begin with, a correlated equilibrium is defined as follows. Let 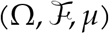 be a finite probability space. Let 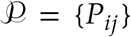 be a collection of pairwise disjoint sets in 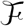 such that ∪_*j,k—*_*P*_*ij*_ = Ω. Let *σ* : Ω → *S* and *σ′* : Ω → *S* for players 1 and 2. The payoff to player 1 playing player 2 with event *ω* is written *u*(*σ*(*ω*), (*σ*′(*ω*)). Player 1 obeys the choreographer for signal *i* if, for all *s* ≠ (*σ*(*ω*),

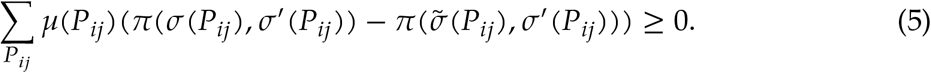

A correlated equilibrium is a state where both players obey all signals.

Given norms *n* = (*L,P,D*) and *n′* = (*L′,P′,D′*) on graph *G* with adjacency matrix *A*, *n* and *n′* form a correlated equilibrium if they are empirically validatable with respect to one another (i.e. Equation 3 holds for both norms, *D* = *P′L′AL*^*T*^, and *D′* = *PLAL′^T^*). Note that Equation 3 is mathematically equivalent to 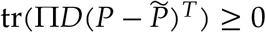 for all 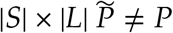. Without loss of generality, let us address player 1. We have 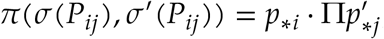, 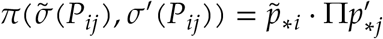, and 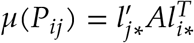. Thus,

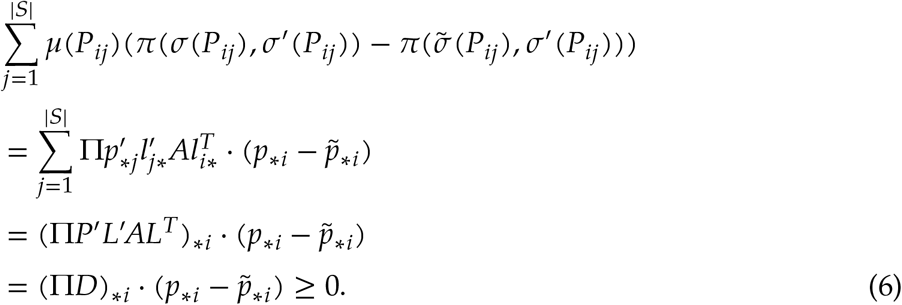

## Appendix B Definitions of norms

We have developed several classes of social norms that are pertinent for evolutionary and stability perspectives. The relations to one another can be seen in Figure 2, and a summary in Table 1.

We define evolutionary stability of a norm as a state in which a few mutants cannot invade [35]. Thus, mathematically, a social norm *n* is an evolutionary stable norm if for every other norm *n′*

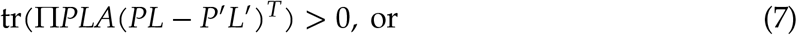

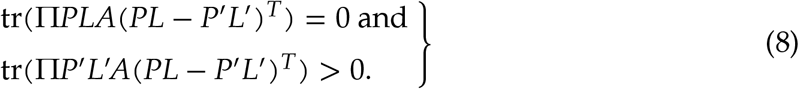

## Appendix C An example of a non-dominant single Nash

We have primarily been concerned with games in which there are multiple pure Nash equilibria in our examples. Additionally, our model has little effect on the Prisoner’s Dilemma where there is a single pure Nash equilibrium. Since, the null norm and all rational norms recommend defection at every vertex. However, this is not true in general for games with a single pure Nash Equilibrium. Such games may have more complicated correlated equilibria beyond that of the pure Nash. For example, consider the Rock-Paper-Scissors game with the addition of a fourth strategy, Thumb, with the following payoff matrix

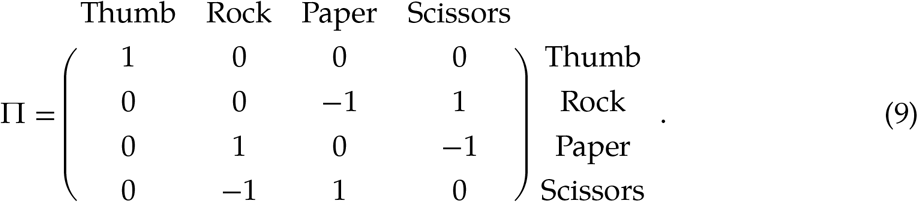

The only pure Nash equilibrium is to play Thumbs. However, there is a mixed Nash equilibrium (and thus also a correlated equilibrium) of playing Rock, Paper, and Scissors with probabilities ⅓ each.

## Appendix D Nash Bargaining Game

Here we detail our model of the Nash Bargaining Game. We assume an atomic distribution of demands to divide a dollar with refinement 1/*α* (*α* > 2, *α* ∈ ℕ). A strategy at vertex *h*, *s*_*h*_, is thus an *α* + 1 probability vector; the entries of which are the probability to demand *i/α* of the dollar. The expected payoff of player *m* vs *n* at edge {*υ*_*h*_,*υ*_*k*_} with 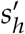 and *s*_*k*_, respectively, is

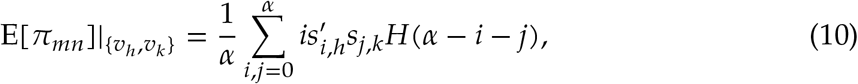

where *H* is the Heaviside function with *H*(0) = 1. The mixed Nash equilibrium strategy is

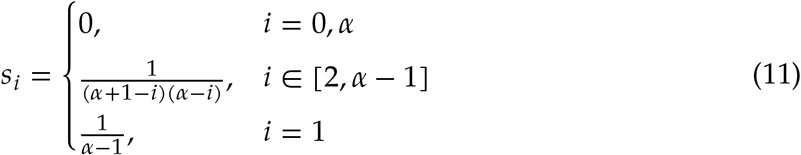

with E[*π*_*nn*_]|_{*υ*_*h*_,*υ*_*k*_}_ = 1/*α*.

We conducted trait substitution simulations with this model. Initial conditions were the single null norm playing the above mixed Nash and a single label applied to all vertices. The algorithm worked as follows: a norm, *n*, is chosen and one of its traits, label or strategy prescription, was randomly chosen and mutated to produced norm *n′*. A prescription was changed with probability 0.5, a label was changed to another currently in use by the norm with probability 0.25 if |*L*| > 1. Otherwise, a label was changed to a random label. The total number of labels available was |*V*|. The descriptive matrix was also changed such that the prescription would be obeyed. If the mutation was neutral, *n′* replaced *n*. If *n′* could invade, we ran the replicator equation for the system with initial conditions *x*_*n*,0_ = max(*x*_n_ − 0.05,0) and *x*_*n*′,0_ = min(*x*_*n′*_ 0.05). At *t* = 100, any norm below the threshold 0.05 is discarded and the remaining population normalized. We plot the results for the complete, ring with degree 2*K*, and small world networks after 1000 iterations of the algorithm. The small world network was constructed via the Watts-Strogatz algorithm with the following initial conditions: a ring graphs with degree 2*K*_0_, and *β* = 0.2 (where β is the rewiring rate).

For completion, Figure 5 depicts the results for the ring graph for *K* = 2 and small world networks generated from ring graphs with *K*_0_ = 1,2. The small world graphs produce similar results to the ring graph. Increasing *K* and *K*_0_ decreases variance when *α* = 4. Further, we view fewer unfair stable monomorphic states as we increase the number of vertices.

**Figure 5:**
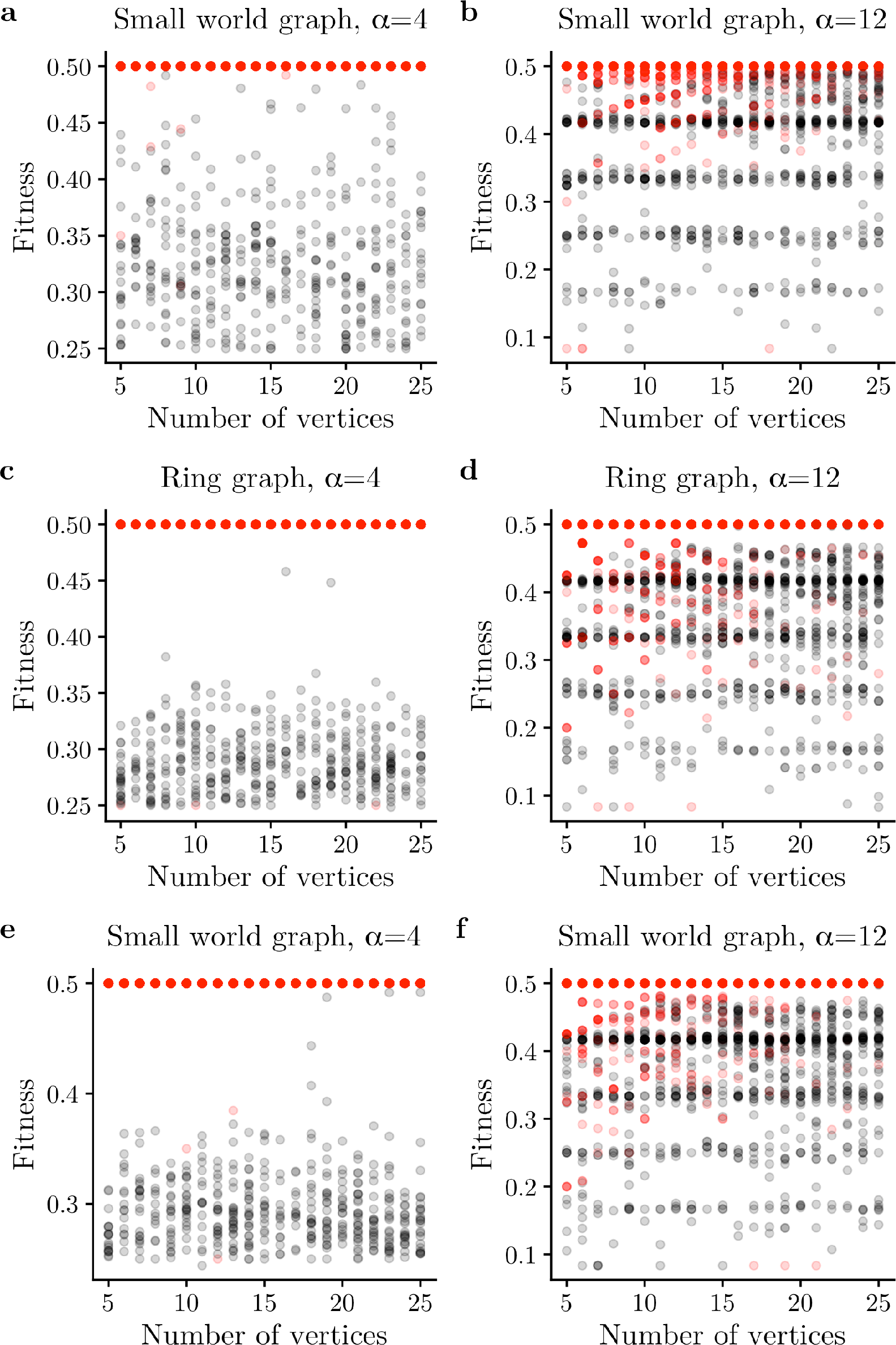
Mean fitness frequency in the Nash Bargaining Game for the ring and small world graphs, and for dollar refinements *α* = 4,12. *K*_0_ = 1 for **a** and **b**, and *K,K*_0_ = 2 for **c-f**. Red corresponds to monomorphic states, and black polymorphic.

## Appendix E Code availability

Data were generated via simulations written in Julia. The code is available at https://github.com/erolakcay/BlindChoreographer.

## Acknowledgements

Funding was provided by Defense Advanced Research Projects Agency NGS2 program (Grant D17AC00005) and the Army Research Office (W911NF-12-R-0012-03).

